# Common object representations for visual production and recognition

**DOI:** 10.1101/097840

**Authors:** Judith E. Fan, Daniel L. K. Yamins, Nicholas B. Turk-Browne

## Abstract

Production and comprehension have long been viewed as inseparable components of language. The study of vision, by contrast, has centered almost exclusively on comprehension. Here we investigate drawing — the most basic form of visual production. How do we convey concepts in visual form, and how does refining this skill, in turn, affect recognition? We developed an online platform for collecting large amounts of drawing and recognition data, and applied a deep convolutional neural network model of visual cortex trained only on natural images to explore the hypothesis that drawing recruits the same abstract feature representations that support natural visual object recognition. Consistent with this hypothesis, higher layers of this model captured the abstract features of both drawings and natural images most important for recognition, and people learning to produce more recognizable drawings of objects exhibited enhanced recognition of those objects. These findings could explain why drawing is so effective for communicating visual concepts, they suggest novel approaches for evaluating and refining conceptual knowledge, and they highlight the potential of deep networks for understanding human learning.

---

Since the earliest known etchings onto cave walls were made 40,000-60,000 years ago in modern-day Spain (Hoffmann et al., 2018; Pike et al., 2012) and Indonesia (Aubert et al., 2014), people have devised many ways to render their thoughts in visual form, employing media ranging from stone and clay to paper and digital displays. The most basic and direct among these visualization techniques is drawing, in which a person produces marks that form an image. Drawing constitutes just one of several methods for externalizing mental representations in graphical form, an ability we term *visual production*. Drawn images predate symbolic writing systems (Clottes, 2008), are pervasive in many cultures (Gombrich, 1989), and are produced prolifically by children from an early age (Kellogg, 1969). Moreover, drawing is a versatile medium supporting diverse representational goals, including photorealistic rendering from observation (Cohen & Bennett, 1997; Tchalenko, 2009), informal sketching from semantic knowledge (Bozeat et al., 2003; Karmiloff-Smith, 1990), and the production of schematic diagrams to support abstract reasoning (Bauer & Johnson-Laird, 1993; Novak & Bulko, 1992)

Here we ask how sketches communicate abstract information about object identity. Remarkably, we perceive simplified sketches of objects as resembling physical objects (Gibson, 1971; Biederman & Ju, 1988; Eitz, Hays, & Alexa, 2012) in spite of the fact that sketches and objects are profoundly different in composition. What operations performed by the brain allow us to see this correspondence in a single glance? Because drawings are composed of contours, it is intuitively appealing to think of a sketch as approximating the edge contours in an image of an object. And because edges can be extracted by applying relatively shallow computations on images, one might expect this resemblance relation to be captured early in visual processing (Hubel & Wiesel, 1968; Marr & Hildreth, 1980; Kay, Naselaris, Prenger, & Gallant, 2008). However, while this might explain why a tracing of an object in a photo might resemble that object (Ishai, Ungerleider, Martin, & Haxby, 2000; Biederman & Ju, 1988), it does not explain humans’ robust ability to recognize the referent of most sketches produced by non-experts, which omit many details, distort the size and proportions of constituent parts, and are highly schematized (Sangkloy, Burnell, Ham, & Hays, 2016; Eitz et al., 2012).

On the other hand, several influential theorists have argued that socially-mediated experience with pictorial representations is necessary to be able to understand them. On this view, generic visual computations are insufficient to account for the perceptual correspondence between drawings and objects — rather, drawings come to denote objects through culturally-specific conventions much the same way that verbal labels denote objects (Goodman, 1976; Gombrich, 1969). While a strong version of this view is controversial (Kennedy, 1975; Gibson, 1971; Abell, 2009), the interpretation of drawings *can* crucially depend on social and cultural factors. Indeed, many drawings that people use to communicate are quite sparse, emphasizing information that is currently relevant, and omitting other details. In the appropriate context, even a few strokes can express the identity of a face (Bergmann, Dale, & Lupyan, 2013), a suggested route (Agrawala & Stolte, 2001), or an intention to act (Galantucci, 2005). Over time, repeated communication between members of the same community can lead to novel graphical conventions (Garrod, Fay, Lee, Oberlander, & MacLeod, 2007; Fay, Garrod, Roberts, & Swoboda, 2010). However, in the absence of contextual cues or prior interaction history, such sparse representations may be insufficient to communicate specific meanings (Healey, Swoboda, Umata, & King, 2007).

How can these different accounts for the basis for fluency with drawn images be reconciled? Here we propose a computational framework for systematically investigating how drawings convey visual concepts. The current investigation is guided by the central hypothesis that the act of producing recognizable drawings of visual objects recruits the same perceptual representation used for recognizing that object in natural scenes. This lays the foundation for future work to understand how drawings that crucially depend on social and cultural context to be recognizable arise from the integration of this common perceptual representation and social learning mechanisms.

A first prediction of our current hypothesis is that drawings of objects — although impoverished in many ways — retain precisely those features that enable recognition of real-world objects. To test this prediction, we employed a deep convolutional neural network model of the ventral visual stream (Yamins et al., 2014; Hong, Yamins, Majaj, & DiCarlo, 2016) to characterize high-level perceptual features of drawings. We compared these feature representations in different layers of the model to those that support the identification of objects in photos. This led to the discovery of a striking isomorphism in the representations of object categories in drawings and photos.

A second prediction is that learning how to draw might refine the representations shared by both drawing and recognition. To test this prediction, we trained people to draw a set of objects and examined, across several experiments, how their drawings of these objects improved and what effect this had on recognition of the objects. To quantify drawing performance, we assessed how well the deep neural network model could recognize the object being drawn. We found that it performed better in classifying drawings after training and that these improved drawings exhibited less feature-level overlap with each other, suggesting that practice drawing these objects had differentiated their underlying representations. This was further reflected in a psychophysical recognition task as enhanced categorization of the objects that people had learned to draw.

These findings provide a first direct demonstration, to our knowledge, that visual production can alter object representations. They also resonate with a substantial literature that has documented a positive relationship between drawing expertise and visual cognition, including tasks tapping perceptual reorganization (Chamberlain, McManus, Riley, Rankin, & Brunswick, 2013; Kozbelt, 2001), encoding of complex object structure into visual memory (Rosenblatt & Winner, 1988; Perdreau & Cavanagh, 2014, 2015), and attentional selection of relevant features to include in depictions (Kozbelt, Seidel, ElBassiouny, Mark, & Owen, 2010; Ostrofsky, Kozbelt, & Seidel, 2012). One limitation of the typical approach taken in these prior studies is that measuring correlations between drawing expertise and visual task performance across individuals cannot provide direct evidence for causal relationships between domains. By experimentally manipulating experience drawing specific objects within-participant and measuring the consequences of this experience on subsequent drawing or recognition of those objects, our study directly tests for such a causal relationship at the granularity of individual objects.

Outside the realm of drawing expertise, figurative drawings have long provided inspiration for scientists investigating the representation of object concepts in early life (Minsky & Papert, 1972; Karmiloff-Smith, 1990). This work has revealed several important insights into the development of children’s ability to include semantically relevant information in their drawings (Sitton & Light, 1992; Light & Simmons, 1983), and to modify their drawings according to the task context (Davis, 1983). Other studies have employed drawing tasks to reveal altered semantic (Bozeat et al., 2003) and spatial (Vallar, 1998) representations in clinical populations. In each of these literatures, a major barrier has been the lack of principled quantitative measures of high-level perceptual information in drawings. As such, previous studies employing drawing tasks have typically relied on qualitative assessments of drawings based on provisional criteria specific to each study, or ad hoc quantitative criteria (Goodenough, 1963), limiting their ability to make detailed predictions on new tasks or datasets.

The approach taken here of using deep convolutional neural network models to characterize the high-level properties of drawings enhances the scientific potential of visual production as a window into human perception and learning. Higher layers of these models both capture adult perceptual judgments of object shape similarity (Kubilius, Bracci, & de Beeck, 2016) and predict neural population responses in categories throughout object-selective cortex (Yamins et al., 2014; Khaligh-Razavi & Kriegeskorte, 2014; Güçlü & van Gerven, 2015). Thus, features learned by these models provide a principled choice of basis for extracting high-level perceptual features from arbitrary visual inputs — including drawings — and measuring changes in their perceptual properties as a consequence of learning. While there is precedent for such model-based analyses of sketches in the machine learning literature (Yu, Yang, Song, Xiang, & Hospedales, 2015; Sangkloy et al., 2016), measuring and explaining *human* learning with these models is innovative and could find broad applicability across many fields, including cognitive development, cognitive neuropsychology, education, communication, and human-computer interaction.

## Generalized object representations

Recognition of visual objects is achieved by a set of hierarchically organized brain regions known as the ventral visual stream (Goodale & Milner, 1992; Rolls, 2000; Malach, Levy, & Hasson, 2002). Simple visual features encoded in lower areas (e.g., orientation, spatial frequency in V1) are successively combined and transformed into more complex features across levels of the hierarchy (Gross, Rocha-Miranda, & Bender, 1972; Hung, Kreiman, Poggio, & DiCarlo, 2005), allowing for read out of abstract object properties from higher areas (e.g., category, identity in inferior temporal [IT] cortex). Recently, these computations have been modeled using deep convolutional neural networks. Such models can approach human-level performance in challenging object recognition tasks and learn features that predict neural population responses in multiple sites along the ventral stream, including V4 and IT (Krizhevsky, Sutskever, & Hinton, 2012; Yamins et al., 2014; Khaligh-Razavi & Kriegeskorte, 2014; Güçlü & van Gerven, 2015). As such, they present an attractive candidate for investigating object recognition when it requires invariance across image domains, such as the recognition of drawings.

We tested the hypothesis that training a deep convolutional neural network to recognize photographs of objects across variable views (Yamins et al., 2014; Hong et al., 2016) would provide a sufficiently robust basis set of high-level features to enable the model to also recognize simple sketches of objects (Fig. 1A). We further predicted that the model’s representations of sketches and photographs should become progressively more similar across successive layers, and peak at the highest layer (approximating IT), consistent with the notion that sketches of objects possess the same abstract features used to recognize natural objects. However, because of large differences between sketches and photographs at the pixel level, we expected their representation in early layers of the model to be much more distinct.

**Figure 1:**
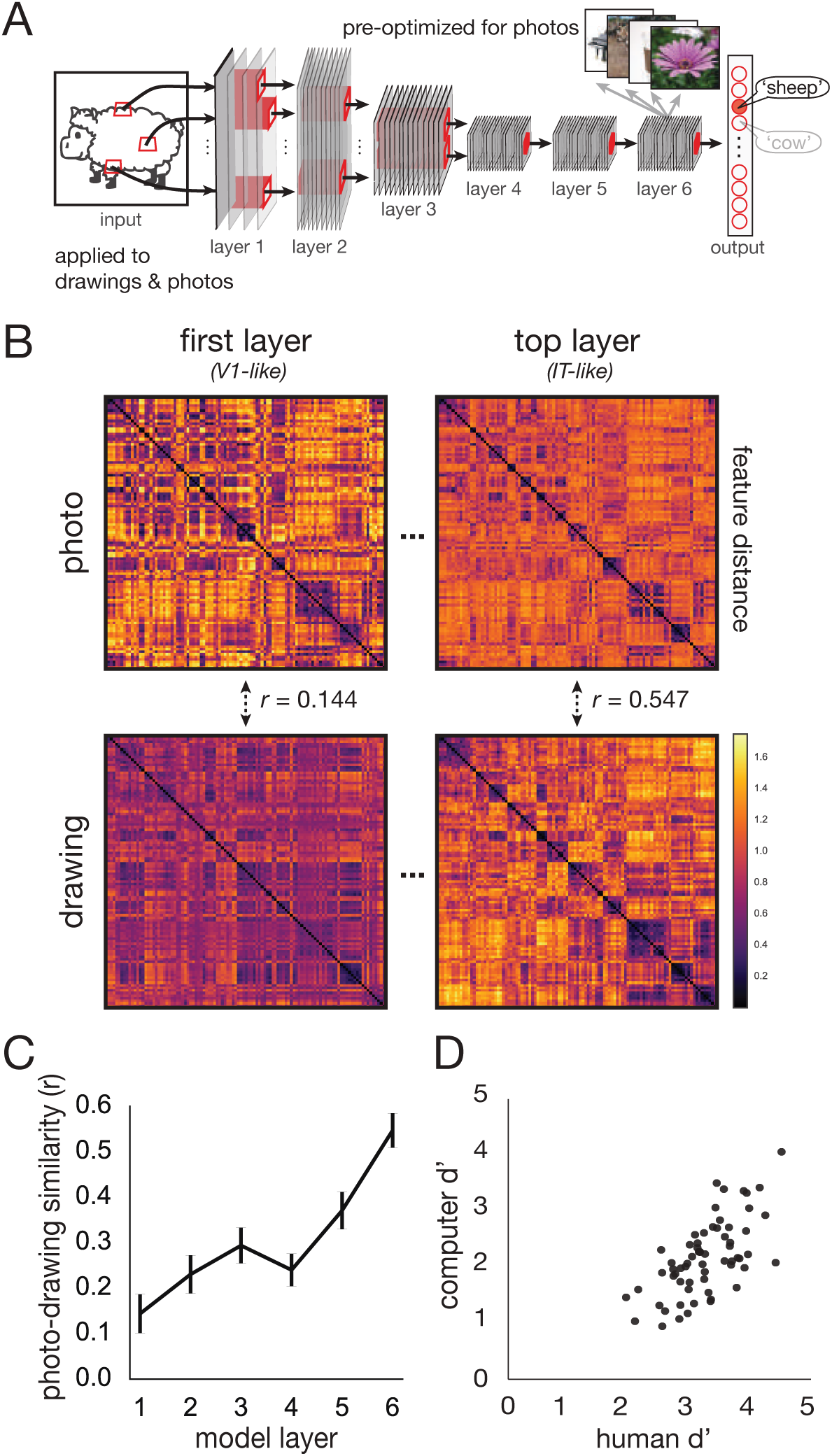
A: Features extracted from drawings and photos using a deep convolutional neural network model optimized to recognize objects in photographs but not drawings. B: Representational distance matrices (RDMs) of model features for each image domain. C: Cross-domain similarity between image domains increases as a function of model layer. Error bars represent 1 s.e.m. D: Human and model recognition performance (d’) was highly consistent across objects (*r* = 0.649, *p* < 0.001.

### Methods

#### Imageset

We obtained 8,400 drawings of 105 common, real-world objects from an existing corpus (Eitz et al., 2012). From the Imagenet database (Deng et al., 2009), we acquired 22,843 photographs of the same 105 objects, depicting diverse exemplars from each object category embedded in natural backgrounds.

#### Computational model

We extracted their features using a deep convolutional neural network model that had been developed using hierarchical modular optimization, a procedure for efficiently searching among mixtures of convolutional neural networks for candidate hierarchical model architectures that achieve high performance on basic-level object recognition (Yamins et al., 2014). This training procedure was performed on an independent image dataset containing millions of photographs from hundreds of object categories other than the 105 categories in our study (Deng et al., 2009). In addition to approaching human-level performance in recognizing these objects, the higher layers of the model quantitatively predict neural population responses in high-level visual cortex (e.g., V4 and IT; Yamins et al., 2014; Hong et al., 2016). As such, this model was an attractive candidate for investigating object recognition invariant to image domain.

Two generations of the hierarchical convolutional neural network model were employed here. The first-generation model (Yamins et al., 2014) was used for the initial experiments, including to extract features from drawings and photographs for the representational distance analyses (Fig. 1) and to provide object-label feedback to participants in the visual production experiment (Fig. 2). The second-generation model (Hong et al., 2016), which became available partway through the study, was used for all subsequent feature analyses, owing to its superior performance, resulting from adding error backpropagation to the training of filter weights.

**Figure 2:**
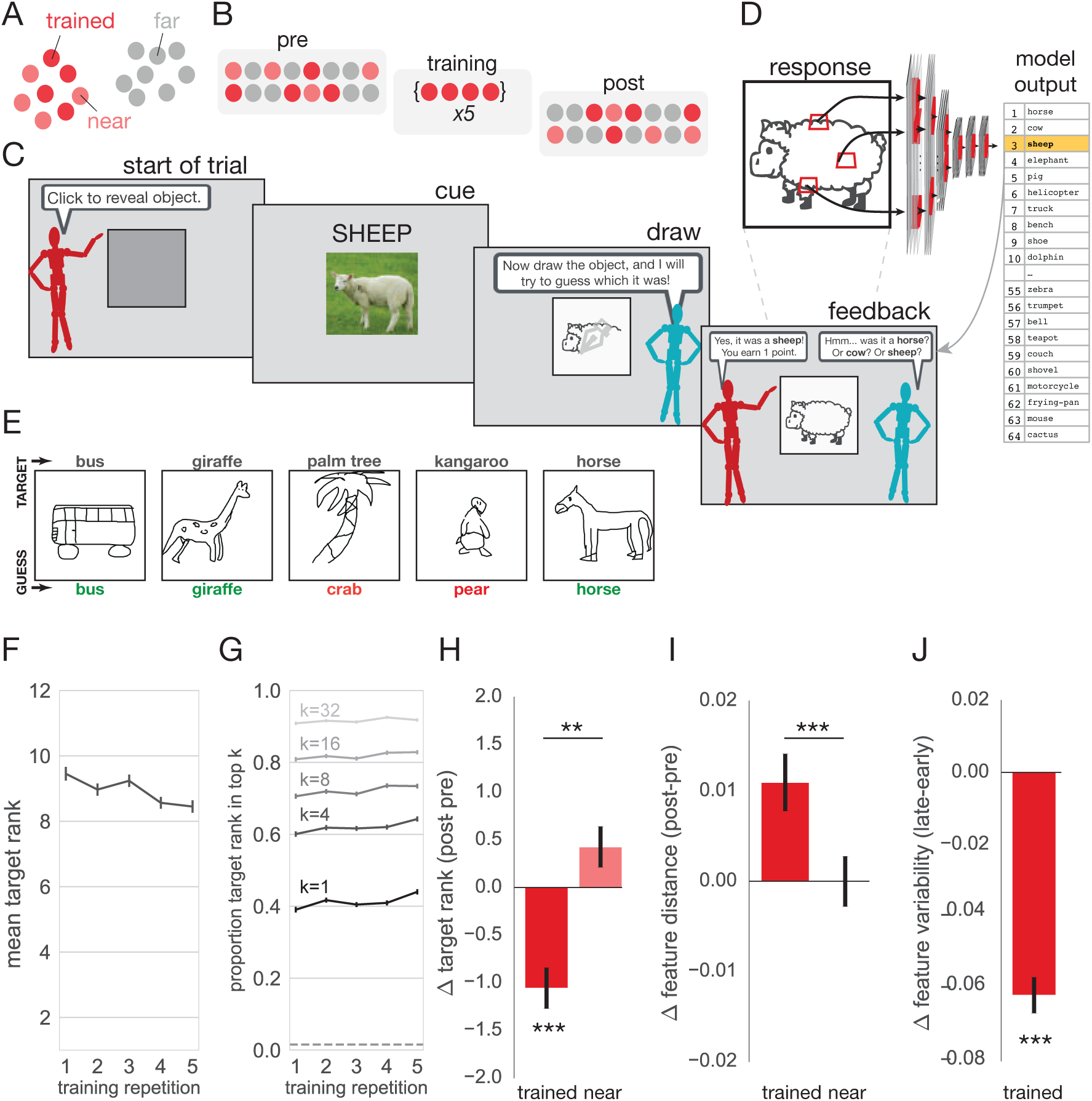
A: Participants randomly assigned two object categories. B: During training, participants repeatedly drew four objects in one category (Trained). Before and after training, participants drew the remaining four objects in that category (Near), and the objects in the second category (Far). C: On each trial, participants were cued to draw an object. D: The neural network model guessed the identity of the drawn object in real time. The rank of the cued object in the ordered list of guesses was used to track changes in performance. E: Sample drawings correctly classified by the model and examples that induced model confusion. F: Mean target rank by training repetition number. G: Proportion of trials on which target rank was within the top *k* labels, by training repetition number. H: Change in performance for Trained and Near vs. control Far objects. I: Change in mean feature distance between objects in Trained and Near vs. Far conditions. J: Change in root-mean-squared feature distances among early (first-three) and late drawings (final-three) of Trained objects. *\*p* < 0.05. *\*\*p* < 0.01. *\*\*\*p* < 0.001. Error bars represent within-participant s.e.m.

#### Representational distance analysis

Each image produces a pattern of feature activations at every layer in the model, each pattern being equivalent to a vector in a feature space with the same number of dimensions as units in that layer. Separately for the drawing and photograph domains, we averaged the feature vectors within each object class for a given layer. To evaluate similarity between domains at each layer, we computed both the Pearson correlation distance (1-*r*) between feature vectors for corresponding classes in each domain. We also computed the matrix of Pearson correlation distances between class vectors within-domain (Kriegeskorte et al., 2008). Formally, this entailed computing: 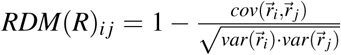, where 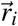 and 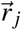 are the mean feature vectors for the *i*th and *j*th object classes, respectively. Each of these 105×105 representational dissimilarity matrices (RDMs) provides a compact description of the layout of objects in the high-dimensional feature space inherent to each layer of the model (Fig. 1B). Following Kriegeskorte et al. (2008), we measured the similarity between object representations in different layers by computing the Spearman rank correlations between the upper triangle of RDMs for those corresponding layers.

Estimates of standard error for the Spearman correlation between RDMs (i.e., between domains or between layers) were generated by jackknife resampling of the 105 object classes. This entails iterating through each of the 105 subsamples that exclude a single class, computing the correlation on each iteration, then aggregating these values. Specifically, the jackknife estimate of the standard error can be computed as: 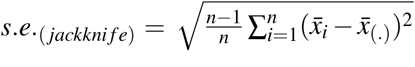, where 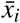 is the correlation based on leaving out the *i*th object class and 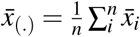, the mean correlation across all subsamples (of size 104). This estimate of standard error allows us to construct 95% confidence intervals and compute two-sided p-values for specific comparisons (Tukey, 1958; Efron, 1979).

#### Classification of drawings

Model features were also used to train linear SVM classifiers (http://scikit-learn.org/) with L2 regularization to evaluate the degree to which category information was linearly accessible in each model layer. Linear classifiers determine a linear weighting of the feature activations which best predicts classification labels on a sample set of training images. Predictions are then made for images held out from the training set, and accuracy is assessed on these held-out images. The robustness of classifier accuracy scores was determined using stratified 5-fold cross validation on 80% train/20% test class-balanced splits.

### Results

Feature vectors corresponding to the same class were more highly correlated than those between classes at all layers (*p* < 0.001), but this correspondence was stronger in the top layer (*r_top_* = 0.223) than in the bottom (*r_bottom_* = 0.072), or any other layer (*p* < 0.001). The same pattern was expressed in the representational dissimilarity matrices (RDMs), with matrices from the top layer exhibiting high similarity (*r* = 0.547, *p* < 0.001), and those from the bottom exhibiting low similarity (*r* = 0.144, *p* < 0.001). Indeed, similarity increased over successive layers in the model (Spearman correlation, *r* = 0.943 *p* < 0.001; Fig. 1C). Consistent with the representational distance analyses, a 105-way support vector machine (SVM) linear classifier was highly accurate for the top layer (64.8% vs. chance= 0.95%, s.e.m. = 1.1% across 5 cross-validation splits), and substantially more so than in the bottom layer (18.6%, cross-validated, s.e.m. = 0.6%).

The model was trained to identify photographs based on human-provided labels, so we interpret this successful recognition of drawings as mirroring how humans would recognize the drawings. To validate this assumption, we recruited an independent cohort of human participants (N=327) to provide labels for drawings from a subset of these categories. As expected, human and computer recognition performance (d’) was highly consistent across objects (*r* = 0.649, *p* < 0.001; Fig. 1D). Thus, the model’s pattern of correct identifications and confusion errors was similar to that of humans performing the same task.

Together, these results show that building a computational model that achieves the level of visual abstraction required to recognize real-world objects under high image variation yields converging feature representations for object drawings and photographs. In other words, the high-level features that support natural object recognition are captured in drawings, allowing a model embodying these features to easily generalize to artificial line drawings. This is consistent with our broader hypothesis that the abstract representation of an object formed during recognition may provide the basis for producing a recognizable image of the object by drawing. A further implication of these results is that drawings may be so effective at conveying visual concepts in part because they take advantage of computational mechanisms already in place (i.e., along the ventral stream) to extract abstract information, such as object identity, from natural visual input.

## Visual production training

If drawing an object involves accessing the representation used to recognize it, then learning how to draw the object better may refine this representation and improve recognition. Here we test for improved recognition in the deep neural network model; in the next section we test for improved recognition in humans. We hypothesized that training people to draw objects would enhance the model’s ability to recognize their drawings of these objects, and that this occurs because they learn to emphasize those features of an object that distinguish it from other objects. This makes the specific prediction that the model’s top-layer representations of the trained objects should differentiate from each other. Such differentiation has traditionally been induced using recognition tasks (Goldstone, 1998), but here we examine it as a consequence of training in visual production.

### Methods

#### Stimuli

A natural starting point for examining learning is to identify objects for which untrained participants have trouble producing recognizable drawings — that is, objects whose drawings are frequently confused by the model with drawings of other objects. In order to identify groups of objects that are drawn similarly prior to training, we applied a clustering algorithm (affinity propagation with damping = 0.9; (Frey & Dueck, 2007)) to the model’s top-layer feature representation of drawings from the same dataset analyzed above (Eitz et al., 2012). This yielded 16 clusters containing between 3 and 20 objects each. Among clusters containing at least 8 objects, we defined 8 “visual categories” containing 8 objects each (Table 1). Each participant was randomly assigned two of these categories, and only the 16 objects from these two categories appeared as drawing targets during their session.

**Table 1:** Objects belonged to eight visual categories, each containing eight items. These categories were derived by applying a clustering procedure to the high-level feature representation of drawings from the Eitz et al. (2012) corpus. Mutual confusion rate reflects the percentage of human labeling errors that involved another object belonging to the same category as the target object (uniform = 11%).

| Category | Objects | Mutual Confusion Rate |
| --- | --- | --- |
| 1 | airplane, blimp, crocodile, fish, helicopter, ship, trumpet, violin | 24.4% |
| 2 | bed, bench, chair, couch, harp, ladder, laptop, table | 59.0% |
| 3 | bell, frying pan, hat, pear, shoe, socks, table lamp, teapot | 29.3% |
| 4 | cat, cow, elephant, horse, kangaroo, pig, rabbit, sheep | 55.7% |
| 5 | banana, dolphin, duck, mosquito, mouse, seagull, shark, swan | 41.0% |
| 6 | floor lamp, fork, guitar, hammer, microphone, shovel, snake, spoon | 37.1% |
| 7 | cactus, crab, giraffe, lobster, palm tree, pineapple, tiger, zebra | 39.2% |
| 8 | SUV, bicycle, bus, motorbike, race car, train, truck, van | 77.3% |

#### Participants

A total of 593 unique participants, who were recruited via Amazon Mechanical Turk (AMT), completed the experiment. Owing to the novelty of the paradigm employed in this study, we could not rely upon preexisting studies to estimate effect sizes. Instead, we used our experimental design as a guide to develop a target sample size that included a few participants (i.e., at least 5) for each of the 56 possible assignments of condition (Trained, Near, and Far) to pairs of categories (out of eight total), and for each of the image-cue and verbal-cue task conditions (see below). Participants were paid a base amount of $1.50 and up to a $3.00 bonus for high task performance. In this and all subsequent studies, participants provided informed consent in accordance with the Princeton IRB. Allocation of participants to groups was conducted anonymously via AMT, and thus the investigators were effectively blinded to the assignment of participants to group and condition during data collection.

#### Experimental procedure

Training was conducted via an online game we developed (“Guess My Sketch”), in which participants (N=593) were repeatedly cued by one avatar to communicate particular object concepts to another avatar representing the model by producing drawings (Fig. 2C). Participants initiated each trial by clicking a central gray square (500 × 500 pixels). Then, a red avatar cued participants to draw an object. Approximately half of participants were cued with a verbal label (N=282) and the other half with a trial-unique photo (N=311). In the image-cue condition, each trial used a unique photograph of the target object on a natural background and participants were instructed to “make a drawing in which someone else is likely to recognize the object depicted” but were informed that the drawing did not have to exactly depict what was in the photo. In the verbal-cue condition, the label of the target object appeared below the square. After cue offset, the blue avatar appeared, prompting the participant to begin drawing. Drawing responses were collected on a digital canvas (500 × 500 pixels) embedded in a web browser window using Raphael Sketchpad (https://ianli.com/sketchpad). Participants drew in black ink (pen width = 5 pixels) using the mouse cursor, and were not able to delete previous strokes. There were no restrictions on how long participants could take to make their drawings, and on average they spent 30.7 s per drawing (95% confidence interval = 8 - 86s).

When the participant’s drawing was submitted on each trial, the blue avatar listed its top three guesses as to the identity of the cued object, thus providing participants with immediate feedback about the recognizability of their drawing. Participants earned points if any of these guesses were correct, proportional to its position in the top three. These guesses were generated by submitting the drawing bitmap to the model running on a server in real-time and passing the top-layer responses though a 64-way support vector machine (SVM) linear classifier pre-trained on photos of the objects from all categories used in this study. The classifier returned a list of 64 margin values, corresponding to the level of confidence that the test image belonged to each object class (Fig. 2D, E).

The rank of the cued object in this list provided a measure of the goodness-of-fit of the submitted drawing to the cued object’s representation in the model and thus served as our primary measure of drawing recognizability (lower rank means better performance). By mapping raw classifier margin values to the rank scale, this ensures that scores for all drawings fall in the same interval, enabling straightforward comparisons across repetitions, objects, and participants. Nevertheless, even when we conduct our main analyses on the raw margins, we find a similar pattern of results. The three objects with highest rank were returned to the participant as guesses. In the verbal-cue condition, when none of the top three guesses were correct, the rank of the target object in this ordered margin list was also returned to the participant (e.g., “Too bad…‘giraffe’ would have been my 9th guess.”).

During the training phase, participants drew four randomly selected objects in one category (Trained) multiple times. Before and after training (Fig. 2B), participants produced one drawing of each of those objects, of each of the other four objects in that category (Near), and of all of the objects in the second category (Far). These conditions allow us to assess the specificity of training effects: Trained objects provide a measure of object-specific learning, Near objects provide a measure of category-specific learning, and Far objects provide a baseline measure of generic task-level or motor improvement. We hypothesized that the Trained objects would become more recognizable to the model after training, relative to both their recognizability before training and to the recognizability of the Near and Far objects.

#### Model validation: category assignments

To validate the model’s representations of the drawings from this experiment, we extracted their features from the top layer. Separately for the image-cue and verbal-cue conditions, we computed the average feature vectors for all drawings of the same object and computed correlation matrices of these average vectors. In both cue conditions, the objects within each of the 8 categories were more similar to each other (image-cue: *r* = 0.633, verbal-cue: *r* = 0.623) than to objects in other categories (image-cue: *r* = −0.083, verbal-cue: *r* = −0.072; *p*s<0.001 based on object-level resampling). The matrices from the two cue conditions were also highly similar to each other (Spearman’s *r* = 0.897), showing that the model successfully captured object identity in both task settings. Moreover, both matrices were highly similar to that computed based on top-layer features of drawings of these 64 objects from the Eitz et al. (2012) drawing corpus (image-cue/original: *r* = 0.789; verbal-cue/original: *r* = 0.818).

#### Model validation: classification of drawings

Across all participants and trials, the model achieved top-1 classification accuracy of 39.8%, top-4 accuracy of 60.5%, and top-8 accuracy of 71.0%, well above chance (1.6%). Relatively lower classification performance compared to that obtained on the original sketch corpus by (Eitz et al., 2012) may not be surprising, since we deliberately designed the stimulus set to include objects that would be more frequently confused with one another, within category. Moreover, given that we were primarily interested in measuring relative *changes* in model recognition of participants’ drawings across different training conditions, this was not a major source of concern.

#### Model validation: measuring human drawing recognition

To validate object-label feedback from the classifier, an independent cohort of human participants (N=327) provided labels each drawing from the image-cue condition from the set of 64 object labels. We found that human and model discrimination performance (d’) were highly correlated across objects (Spearman’s *r* = 0.649, Fig. 1D). We additionally computed the mutual confusion rate, defined as the percentage of (first-guess) labeling errors that involved another object belonging to the same category as the target object. If these errors were spread uniformly over all distractors, the expected mutual confusion rate would be: 7/63 = 0.11. All categories exhibited mutual confusion rates reliably above uniform responding (across objects within category: *t*_7_*s >* 2.76, *p*s *<* 0.028), which further validates category assignments. Overall human identification accuracy was 70.2%, which unsurprisingly exceeded model top-1 classification accuracy, and may be explained by the vast amount of additional experience adult human observers have had with a wider variety of visual inputs and tasks than the model.

#### Statistics

Before performing statistical tests, we visualized data and examined assumptions. Quantile-quantile plots revealed a reasonable approximation to normality, an assumption of the paired t-test. Mauchly’s test of sphericity indicated that the assumption of sphericity for the repeated-measures ANOVAs had not been violated. All p-values reported are two-sided. We also found that employing non-parametric analysis techniques (i.e., bootstrap resampling) gave similar results, suggesting that the choice of test (and assumptions therein) does not impact our conclusions. We did not have a prespecified way of handling outliers in this study, so we report analyses with all data included.

#### Results

Because objects were randomly assigned to condition across participants, we expected no differences in mean rank for Trained, Near, and Far objects in the pretest. Indeed, a 3 (condition: Trained, Near, Far) × 2 (cue type: word, image) repeated-measures ANOVA revealed no main effect of condition (*F*(2,587) = 1.16, *p* = 0.315). There was a main effect of cue type (*F*(2,587) = 14.8, *p* < 0.001), with image-cue performance (M = 8.69, SD = 6.55) exceeding word-cue performance (M = 10.24, SD = 7.58). While we did not have strong *a priori* hypotheses about the effect of cue type, one possibility is that image cues may have led to better performance by reminding participants of diagnostic visual details that may be more difficult to retrieve from semantic memory based on the verbal cue alone (e.g., spatial relationships between parts of a bicycle). Regardless, cue type did not interact with condition, as expected (*F*(2,587)=0.147, *p* = 0.863).

During the training phase, participants drew the four Trained objects five times each, in a randomly interleaved order. To assess changes in performance over training, we computed the mean rank across objects for each repetition and assessed its relationship to repetition number. Although there was large variation across participants, the trend across repetitions was reliably negative (mean Spearman’s *r* = −0.075; *t* = 3.37, *p* = 0.0007), demonstrating that drawings improved with practice (Fig. 2F). In order to better understand what was driving these changes in mean target rank, next we computed the proportion of trials on which the target rank was within the top-*k* values, for *k* ∈ {1,4,8,16,32}, across repetitions. Visualizing this timeseries revealed the largest gains for lower values of *k*, indicating that much of the improvement could be explained by the target rank value moving into the top few positions (Fig. 2G).

To assess learning across conditions, we compared differences between pretest and posttest performance (Near and Far objects only appeared during these phases). For each object in all conditions, we calculated the change in rank (Δ*_rank_* = *rank_post_ – rank_pre_*), then averaged these Δ*_rank_* values across objects in each condition. We then performed the same type of ANOVA across participants as for the pretest analysis, revealing a significant difference in rank change between conditions (*F*(2, 1182) = 7.67, *p* < 0.001). There was no main effect of cue type (*F*(1, 591)=1.66, *p* = 0.198), nor an interaction between condition and cue type (*F*(2,1182) = 0.162, *p* = 0.851), so we collapse across cue type in subsequent analyses. Drawings of Trained objects were better recognized by the model following training (Δ*_rank_* < 0: *t*(592) = 4.04, *p* < 0.001, two-sided in this and all subsequent t-tests); no such improvement was found for Near objects (*t*(592) = 0.511, *p* = 0.609) or Far objects (*t*(592) = 1.22, *p* = 0.223), suggesting that training had not led to general improvements at the category level or task level. The improvement for Trained exceeded that of Near (*t*(592) = 3.44, *p* < 0.001) and Far (*t*(592) = 2.91, *p* = 0.004), which themselves did not differ (*t*(592) = 1.15, *p* = 0.252). Taken together, these results indicate that production training primarily resulted in object-specific benefits (Fig. 2H). In particular, improved classification performance for Trained objects was driven primarily by an increase in the hit rate (+5.2%, percentage of trials, pooling across participants; *p* < 0.001, bootstrap resampling of participants), with marginally significant decreases in the rate that Trained drawings were misclassified as being Near (−1.3%, *p* = 0.080) or Far (−1.2%, *p* = 0.082) objects.

The improved rank score for Trained objects shows that their high-level feature representations became more linearly discriminable. We investigated two potential sources of this differentiation (not mutually exclusive): increased feature distances among Trained objects (“separation”) or decreased feature variance of individual Trained objects (“sharpening”).

To test for separation, we first extracted the model’s top-layer feature representation of all drawings from the pretest and posttest and computed a matrix of the correlation distances between these feature vectors (Fig. 2I). Then, for each Trained object, we compared its distance before training with the other Trained objects before training (pre/pre) with the other Trained objects after training (pre/post). The same difference was calculated for Near and Far objects as controls. Increased distance for pre/post vs. pre/pre in the Trained condition relative to the Near and Far baselines would indicate that drawing induced separation of object representations. Consistent with this possibility, a one-way repeated-measures ANOVA revealed a main effect of condition (*F*(2, 1773) = 3.12, *p* = 0.044). Planned comparisons confirmed that Trained objects separated more than Near objects (*t*(592 = 3.48, *p* = 0.0005) and Far objects (*t*(592) = 3.46, *p* = 0.0005), which did not differ from each other (*t*(592) = 0.005, *p* = 0.996).

To test for sharpening, we tracked changes in the distance between feature vectors from the top layer across successive drawings of the same object during training. For each Trained object, we constructed a distance matrix relating drawings across all repetitions (Fig. 2J); this analysis was not possible for Near or Far objects because they were only drawn at the start and end of the study. We quantified change in feature variability in two ways: by comparing root-mean-squared feature distances among early drawings (first three) and late drawings (final three), and by measuring the trend in feature distances across pairs of successive drawings. We found that late drawings were reliably more similar to one another than early drawings (mean Δ =−0.061, s.e.m. = 0.005, *p* < 0.001, bootstrap resampling of participants), and that this reflected a gradual decrease in the amount by which successive drawings differed across repetitions (Spearman’s *r* =−0.211, s.e.m. = 0.022, *p* < 0.001, bootstrap resampling of participants).

To evaluate the respective contributions of separation and sharpening to the improvement in drawing performance (as quantified by the model), we regressed both of these measures on the change in rank for Trained vs. Far objects. Across participants, we found that sharpening (β=7.62, *t*(590) = 2.53, *p* = 0.012) but not separation (β=0.02, *t*(590) = 0.004, *p* = 0.997) predicted model performance. This suggests that decreased variability was most directly responsible for the increased discriminability of Trained objects.

Taken together, these results are consistent with the notion that people had learned through practice which features were most diagnostic of the Trained objects, leading to more consistent expression of these distinguishing features in their drawings over time. While this study cannot disentangle the contribution of task practice from the contribution of semantic feedback from the model to this learning, further insight could be gained from experiments which do separately manipulate task practice, as well as the availability and type of feedback.

## Controlling for visual exposure to drawings

The results so far have been interpreted as a consequence of repeatedly drawing the Trained objects. However, as these objects were being drawn, participants also received additional perceptual experience with drawings of them. This additional visual experience with drawings of the Trained objects suggests an alternative explanation for the benefit for Trained over Near and Far objects, which were only encountered in the pretest and posttest. To evaluate this alternative, we conducted two additional experiments that controlled for the amount of visual exposure to sketches received during training.

### Methods

#### Control experiment 1: viewing finished drawings

For each of the 593 participants in the original cohort, we recruited a new participant to repeat the same sequence of trials. They were paid a base amount of $2.00 and up to a $3.00 bonus for high task performance. However, instead of producing a drawing on each trial of the training phase, they were presented with the same cue provided to their matched participant (i.e., image or verbal cue), then the finished drawing produced by the matched participant. To encourage participants to inspect the sketch, participants had the goal of predicting the *model’s* top guess for the drawing. In some ways, this provides even more perceptual experience, as they always viewed the completed, most recognizable drawing, rather than incomplete, ambiguous versions. Participants typed their response into a text field, and only the labels of the 64 objects in the set were accepted. They had to wait 4000 ms before being able to submit their response. Participants drew all objects once each before and after training, allowing us to measure the consequences of viewing finished drawings on drawing performance.

#### Control experiment 2: observing stroke dynamics

Again, 593 naive participants were recruited via AMT and paid a base amount of $2.00 and up to a $3.00 bonus for high task performance. Again, each participant was matched with one of the original participants and received the same sequence of trials. On each training trial, they were given the same cue provided to their matched participant, then observed a stroke-by-stroke reconstruction of the drawing produced by their matched participant, with a stroked added every 500 ms. They performed the same prediction task as in Control experiment 1 with 4000-ms waiting period from the start of the animation, ensuring that at least eight strokes appeared (or all of the strokes if eight or fewer). Participants drew all objects once each before and after training, allowing us to measure the consequences of observing stroke dynamics on drawing performance.

### Results

We found that viewing finished drawings yielded only modest changes in drawing performance for Trained objects (*t*(592) = 1.72, *p* = 0.086), and no reliable improvement for Trained objects relative to Far (*t*(592) = 0.580, *p* = 0.562) or Near objects (*t*(592) = 1.21, *p* = 0.228). Again, neither Near (*t*(592) = 0.160, *p* = 0.873) nor Far (*t*(592) = 1.27, *p* = 0.205) objects improved relative to their pretest baseline, nor differed significantly from one another (*t*(592) = 0.714, *p* = 0.475). Together, these results suggest that mere exposure to drawings is insufficient for improving the ability to produce recognizable drawings.

Although viewing completed drawings did not improve drawing performance, this only captures part of the perceptual experience of drawing. In particular, completed drawings are the result of composing a series of individual strokes into parts, and parts into whole drawings, over time. We hypothesized that observing these stroke-level dynamics may be more beneficial for learning how to draw, because they convey information about the procedure for composing a drawing that may be subsequently used to support successful drawing.

We found that observing dynamic reconstructions of drawings produced reliable pre-post improvement for Trained objects (*t*(592) = 2.880, *p* = 0.004), which significantly exceeded that of Far (*t*(592) = 2.09, *p* = 0.037) and Near (*t*(592) = 2.55, *p* = 0.011) objects. These findings suggest that observing the process of drawing construction improves participants’ subsequent ability to make recognizable drawings of those objects they had previously observed being drawn. We again found no reliable pre-post changes in performance for Near (*t*(592) = 0.448, *p* = 0.655) or Far (*t*(592) = 0.606, *p* = 0.545) objects, and these conditions did not differ from one another (*t*(592) = 0.751, *p* = 0.453).

In order to evaluate whether learning differed according to the training modality (i.e., drawing vs. viewing finished drawings vs. observing dynamic reconstructions), we compared the degree of improvement for Trained objects across modalities. We found no reliable difference between the drawing and dynamic modalities (*t*(592) = 0.929, *p* = 0.353), and a marginal difference between the drawing and static modalities (*t*(592) = 1.81, *p* = 0.071). However, we caution against strong conclusions about the relative effectiveness of these different training modalities based on these data alone. Rather, exploring how differences in training modality (e.g., production vs. observation) relate to subsequent performance would be a fruitful target for future research.

### Consequences for object recognition

Above we showed that training participants how to draw objects resulted in drawings that were more recognizable to a deep neural network model of the ventral visual stream. The model was pre-trained and its parameters fixed, so these changes in the rank of the cued object can be interpreted as evidence that the object representations of participants were refined by the training. However, although suggestive, this is only indirect evidence for the claim that these refined representations are the ones in the ventral visual stream that subserve human object recognition abilities. Participants may have improved their drawings without any change in their internal visual representation of these objects (i.e., in the ventral stream), for example, as a result of object-specific motor learning of stroke sequences.

To evaluate more directly how learning to draw impacts human recognition we conducted a transfer study (N=72) in which the drawing training phase was bookended by a recognition pretest and posttest. Motivated by the earlier finding that the feature representation of Trained objects in the top, IT-like layer of the model differentiated with training, we hypothesized that drawing would increase the perceptual discriminability between Trained objects. We tested this hypothesis by having participants discriminate morphed versions of these objects (Goldstone, 1998; Livingston, Andrews, & Harnad, 1998), predicting that training would cause morphs between Trained objects to be perceived as more distinct — that is, morphs in the middle of the range should become more consistently recognized as the majority object, resulting in a steeper slope of the psychometric function relating the morphing proportion to object labels. The rationale for this prediction is that differentiation should reduce features shared between object representations such that intermediate morphs are represented more in terms of the distinguishing features of the majority object, supporting more consistent identification of that object.

### Methods

#### Stimuli

The four objects in each of the two categories were selected to allow the construction of a quartet of 3D mesh models (Autodesk Maya) sharing the same vertices, thereby enabling quantitatively precise morphing and full control over category-orthogonal image parameters (e.g., pose, size, background). This resulted in six axes (‘morphlines’) connecting all pairs of objects within each category and 12 total axes for both categories. For each axis, we derived a perceptually uniform space from which to sample morphs, in order to increase sensitivity for measuring the slope of the psychometric function. As a first step, we rendered a series of 12 morphs for each axis, linearly interpolated between the endpoint objects. Each morph was rendered from a 10° viewing angle (i.e., slightly above) at a fixed distance on a gray background in 40 viewpoints (i.e., each rotated by an additional 9° about the vertical axis). We recruited a separate cohort of 40 participants via AMT to provide 288 identification judgments each for random subsets of these morphs, yielding 80 baseline judgments per morph (two per viewpoint). For each morph (e.g., sedan/limo), we computed the proportion of trials that the morph was identified as being one of the endpoint objects (e.g., sedan) and fit these data with a logistic function to derive a population psychometric curve. We used this curve to estimate the morphing levels that produced 0% (or the minimum), 20%, 40%, 60%, 80%, and 100% (or the maximum) identifications as one of the endpoint objects. The resulting six morphs per axis evenly spanned the subjective transition between endpoints and were included in the training study.

**Table 2:**
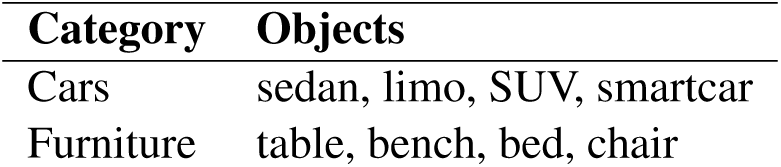
Endpoint objects included in transfer experiments.

#### Participants

Based on initial piloting, we developed a target sample size of 72 participants, across whom all condition and object assignments would be fully counterbalanced. Of the original group of 72 participants recruited from AMT, 25 were excluded because their data could not be fit with a logistic function in at least one condition either before or after drawing training. This occurred either because of non-monotonicity in their psychometric curves (i.e., inconsistent responding) or hypersteepness of the slope parameter (i.e., approaching infinity). We recruited additional participants to fill these sessions to ensure that the design was counterbalanced. In total, 97 participants completed the task and received $5.00.

#### Experimental procedure

We adapted our training task in several ways to enhance our ability to measure the hypothesized perceptual changes. Estimating the parameters of psychometric curves requires many trials, so we used a reduced stimulus set of two categories with four objects each (furniture: table, bench, bed, chair; cars: sedan, limo, SUV, smartcar; Fig. 3A). To enable morphing of these objects, we commissioned an artist to design 3D models of each object with the same number of vertices per category, and then generated intermediate objects via interpolation between these endpoint objects. To increase the probability of inducing transfer effects, we also increased the number of training trials per object from 5 to 16. Finally, we removed feedback from the training phase to more cleanly isolate the consequences of production as such.

**Figure 3:**
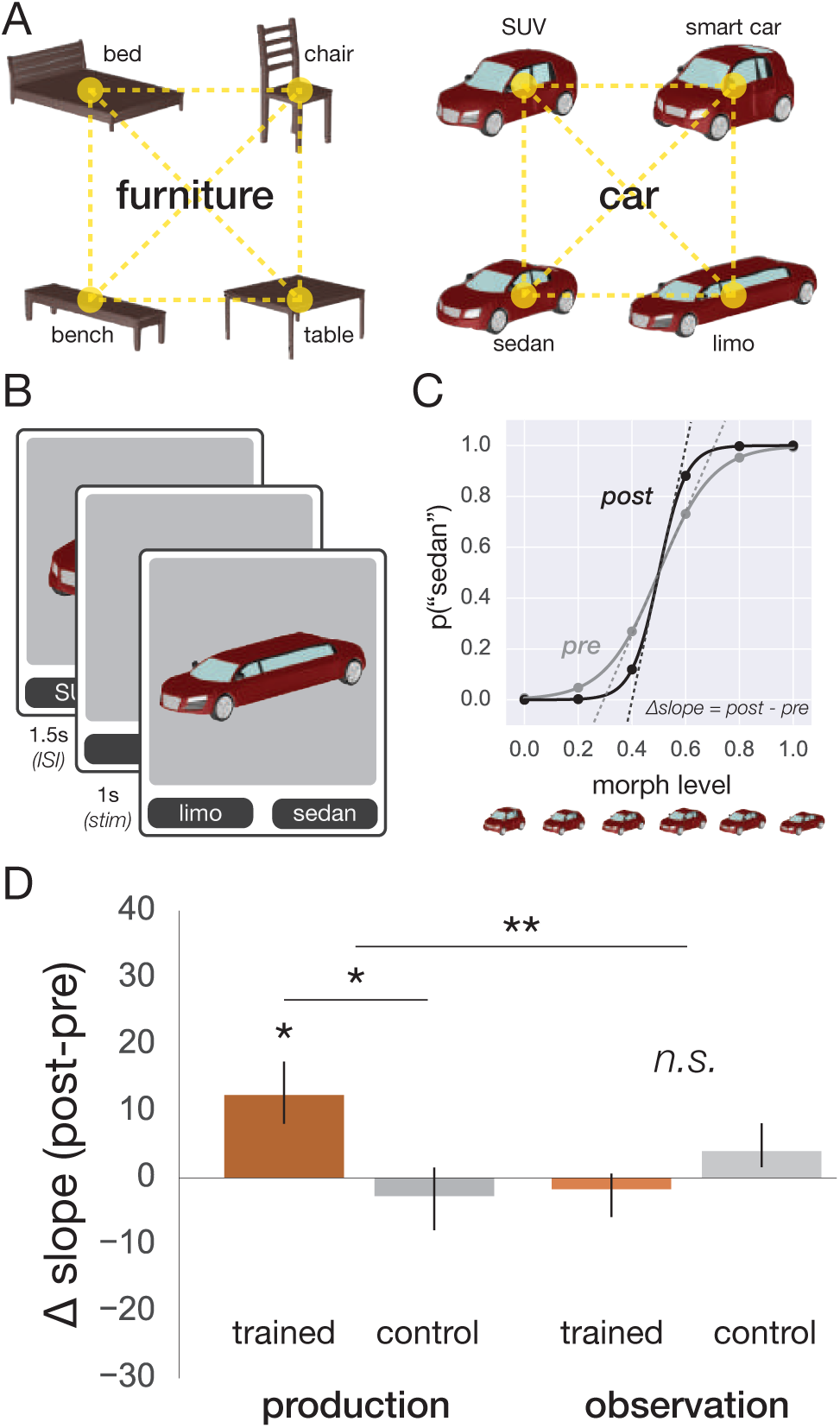
A: Stimuli in the perceptual discrimination experiment. B: Before and after training, participants performed a recognition task in which they discriminated morphs of the two Trained objects, or the two Control objects. C: Psychometric data were fit with a logistic function, whose slope was predicted to increase after training. D: Change in slope for Trained/Control conditions in each group. *\*p* < 0.05. *\*\*p* < 0.01. Error bars represent 1 s.e.m.

Each participant was randomly assigned one of the two categories and was trained to draw two of the objects in that category (Trained). The other two objects (Control) served as a baseline for changes in recognition. At the beginning of each session, participants were familiarized with each of the eight endpoint objects that might appear. On each familiarization trial, an animation of one of the objects continuously rotating was played. Objects appeared on a gray background at the same viewing angle, distance, and viewpoints used in the experiment. The name of the object was displayed in large font above the animation (“This is a SEDAN.”). The participant viewed the object completing at least one full rotation (6s) before proceeding to the next object.

After familiarization, participants completed three phases of the experiment: pretest discrimination, drawing training, and posttest discrimination. On each drawing training trial, participants were cued with one of the Trained endpoint objects (trial-unique viewpoint) and then made a drawing of it. Trained objects were drawn 16 times each during the training phase (32 total trials), in a randomized order and cued with trial-unique viewpoints. No feedback was provided but the task was otherwise identical to the earlier drawing training study.

Before and after training, participants were tested on perceptual discrimination of morphs of the two Trained objects or morphs of the two Control objects. All 12 morphs (six morphs per pair) were shown 12 times each during both the pretest and posttest, always from a trial-unique viewpoint. On each trial, participants were briefly presented (1000 ms) with the morph and made a forced-choice judgment about which of the two objects they saw by clicking one of two labels that appeared below the image (Fig. 3B). The assignment of labels to buttons was randomized across trials. Participants who did not achieve greater than 80% accuracy on the unambiguous 0%/100% endpoint objects in the pretest phase did not proceed to the training phase.

For each participant and for both the pretest and posttest, we constructed psychometric curves for the Trained and Control object pairs, relating different morphing levels to the proportion of trials (out of 12) in which a given object was chosen. These curves were fit with a logistic function 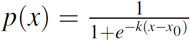 using the Levenberg-Marquardt algorithm as implemented in Scipy (https://www.scipy.org/), where *p*(*x*) is the proportion of trials on which the first object was chosen, *k* represents the slope, and x_0_ the midpoint of the sigmoid. Our key prediction concerned the slope parameter from the fitted logistic curve: if drawing enhances the discriminability of Trained objects, then morphs in the middle of the range to become more consistently recognized as the majority object, and the slope parameter *k* for the Trained pair should increase from pretest to posttest more than for the Control pair (Fig. 3C).

#### Control experiment: observing stroke dynamics

For each of the 72 sessions in the main experiment, we recruited a naive participant to repeat the same sequence of trials, except that during the training phase, they observed a stroke-by-stroke reconstruction of the drawing produced by their matched participant, with a stroked added every 500 ms. Of this initial new cohort of 72 participants, 16 were excluded because their data could not be fit with a logistic function in at least one condition either before or after drawing training. As in the main experiment, we recruited additional participants to fill these sessions to ensure that the design was counterbalanced. In total, 88 participants completed the task and received $5.00. Before and after training, all participants were tested on perceptual discrimination of the Trained and Control object pairs.

#### Statistics

Before performing statistical tests, we visualized data and examined assumptions. Quantile-quantile plots revealed that data did not follow a normal distribution, so classical inference tests (e.g., ANOVA, *t*-test) that rely upon assumptions of normality were not appropriate. Instead, we employed bootstrap resampling (Efron & Tibshirani, 1986) to construct 95% confidence intervals and compute p-values for key parameters and comparisons of interest (i.e., change in slope from pretest to posttest). This entailed resampling 72 participants’ worth of data with replacement, then computing the mean, on each of 10,000 iterations, for each experiment. The two-sided p-value was defined as the proportion of these iterations on which this mean fell below zero, multiplied by two.

### Results

Slope estimates did not differ between Trained and Control pairs during the pretest (*p* = 0.473, bootstrapped resampling), which was expected since there was no difference between conditions prior to training. After training, the slope for the Trained pair reliably increased (*p* = 0.004; Fig. 3D), and more than for the Control pair (*p* = 0.005), whose slope did not change (*p* = 0.533). The threshold parameter did not change significantly for either condition (Trained: *p* = 0.092; Control: *p* = 0.308). These results show that visual production training can generalize to a recognition task, lending key support to our hypothesis that production and recognition engage a common high-level representation for objects.

The question raised earlier about the role of perceptual experience during drawing is especially salient in this study, which employed a perceptual measure. That is, enhanced perceptual discrimination for the Trained pair may reflect perceptual learning due to greater visual exposure to these objects while they were being drawn repeatedly. Although such observation improved drawing performance in the earlier experiment, we hypothesized that transfer to a recognition task would require more significant representational changes, as would be induced by drawing, and thus that this group might not show improved perceptual discrimination. The rationale for this prediction is that drawing produces continuous visual and haptic feedback, and such sensory feedback during visually-guided actions has previously been shown to facilitate transfer to perceptual tasks (Fan, Turk-Browne, & Taylor, 2016).

Indeed, there was no reliable change in slope for either the Trained pair (*p* = 0.475; Fig. 3D) or Control pair (*p* = 0.332), and no difference between these conditions (*p* = 0.169). Moreover, there was an interaction between training group and condition (*p* = 0.002), with a larger increase in slope for Trained vs. Control in the participants who were trained to draw than in those who observed somebody else drawing. The threshold parameter did not change for either condition (Trained: *p* = 0.838; Control: *p* = 0.147). These results suggest that the generalization of drawing training to perceptual discrimination was driven by aspects of visual production beyond observation of the consequences of action.

It is important to note that the current experiment measures explicit categorization performance, thus it is possible that enhanced discrimination of Trained objects reflects changes in how these representations are read out to produce a category label, as opposed to changes in how they are automatically perceived. These possibilities could be disentangled in future studies that rely on short presentation times and a nonverbal response modality. For example, one could test how well participants are able to judge whether two briefly presented morphed objects are the same or different. Faster and more accurate responses on trials where the two morphs straddle the discrimination boundary between objects as opposed to being on the same side of the boundary would provide stronger evidence for automatic effects on perception.

## Discussion

The present study investigated the relationship between the ability to recognize objects — a biological endowment shared with other species – and to produce images of objects by drawing – a relatively recent development from human prehistory. We examined the hypothesis that these two behaviors are at least partly served by a common representational substrate for objects.

We discovered that a deep neural network model trained only on photos succeeded in recognizing drawings, suggesting that this kind of abstraction can arise from the same neural architecture evolved to make sense of natural visual inputs (Sayim & Cavanagh, 2011). These findings argue against a strong version of the hypothesis that line drawings are purely a product of culturally-specific conventions (Goodman, 1976; Gombrich, 1969). Rather, they are consistent with evidence from various domains, including developmental, cross-cultural, and comparative studies of drawing perception. For example, human infants (Hochberg & Brooks, 1962), people living in remote regions without pictorial art traditions and without substantial contact with Western visual media (Kennedy & Ross, 1975), as well as higher non-human primates (Tanaka, 2007) are able to recognize line drawings of familiar objects, even without prior experience with drawings. Moreover, these findings also provide a computational basis for understanding how line drawings can drive object recognition so effectively (Biederman & Ju, 1988; Walther, Chai, Caddigan, Beck, & Li, 2011). We also found that learning how to draw increased the discriminability of trained object representations, as quantified by improved recognition performance and reduced feature overlap in the model and by enhanced discrimination of trained objects in human participants. These findings are reminiscent of the way other generative behaviors such as memory retrieval (Slamecka & Graf, 1978; Crutcher & Healy, 1989; Karpicke & Roediger, 2008) and self-explanation (Chi, De Leeuw, Chiu, & LaVancher, 1994; Williams & Lombrozo, 2013) can powerfully guide learning.

The learning mechanisms responsible for such changes are not yet known, but a promising avenue forward is to build on extant theories of how differentiation between mental representations occurs. Two broad classes of candidate mechanisms may be particularly worthwhile to test: (1) strengthening of diagnostic features of objects through increased weighting of relevant dimensions (Goldstone, 1998), and (2) weakening of features that overlap between objects through competitive dynamics (Norman, Newman, Detre, & Polyn, 2006). These mechanisms are not mutually exclusive, while they do make different predictions about learning outcomes under certain conditions. For instance, in the context of prolonged competition between two similar objects (e.g., alternating drawing of sheep vs. goat), the first mechanism could stabilize and refine object representations in a generalized manner, whereas the second mechanism would exaggerate differences specifically along the axis separating the competing objects in representational space. Our observation that increased recognizability of sketches across training repetitions was accompanied by sharpening of trained object representations, as measured by reduced feature variability within-object, is in principle consistent with both strengthening and weakening mechanisms. In order to tease these two mechanisms apart, a critical variable that future studies might manipulate is the initial similarity between object representations. Under the strengthening mechanism, diagnostic features are predicted to be enhanced regardless of the initial similarity between objects. On the other hand, under the weakening mechanism, diagnostic features that are shared by similar objects are predicted to be weakened, while non-overlapping diagnostic features are still enhanced.

Here we found preliminary evidence for the benefits of observational learning for visual production — that is, observing someone else produce drawings, stroke-by-stroke, led to reliable improvement in the ability to subsequently produce drawings of those same objects (Mattar & Gribble, 2005). This finding raises several important questions regarding the mechanisms responsible for this learning. First, it may be that such drawing demonstrations benefit learning by directing a learner’s attention to a series of smaller curve segments, thereby making it easier to encode and later retrieve an effective sequence of actions to produce a recognizable drawing. This could be tested by having participants view a finished drawing, where each stroke is highlighted in sequence, even in an arbitrary order. Insofar as the decomposition of the drawing into constituent strokes is critical, this form of observational learning should also lead to improvement in visual production. Second, it may be that while observing such demonstrations, learners actively make visual predictions about where strokes will appear and what they will look like. Insofar as the sequence and appearance of strokes in a demonstration deviates from what is expected, this visual prediction error may also promote encoding of the demonstrated sequence, thereby improving subsequent visual production. This could be tested by having participants view the same strokes presented in the reverse (less predictable) or a shuffled (least predictable) order. If the predictability of the stroke sequence modulates learning, moderately unpredictable sequences that generate many small prediction errors might enhance learning, while completely unpredictable sequences may lead to worse subsequent performance. With regards to both the enhanced-attention and prediction-error accounts above, one might further predict that the benefits of such observational learning would be greater when observing higher-scoring drawings, rather than lower-scoring drawings.

We also found that the benefits of observational learning did not generalize to the perceptual discrimination task. There are multiple potential reasons for this that would be valuable to distinguish in future work. One possibility is that immediate sensory feedback from visually-guided action may have a disproportionate effect on visual learning (Fan et al., 2016). If so, then much more observation experience may be required to improve perceptual decision-making to the same degree. This could be addressed in new experiments by varying the length of the observational learning phase relative to the production phase. Another possibility is that the lack of task-related reward feedback in this experiment may have had a particularly strong dampening effect on how effectively participants encoded the observed strokes; this could be addressed in new experiments by employing an engaging cover task during observation that guides attention to each new stroke.

The claim that visual production recruits and refines the same high-level object representation used during visual recognition may appear to be in tension with the distinction between vision-for-action and vision-for-recognition, which are functionally segregated into dorsal and ventral streams, respectively (Goodale & Milner, 1992). However, our findings can be reconciled with this view by considering the type of action investigated here. Namely, our drawing task involves producing a recognizable image of an object held in mind rather than reaching toward or manipulating a physical object in the world. Couched this way, the act of drawing coincides with a core function of the ventral stream – the computation of abstract, geometric properties of objects that are diagnostic of their identity (DiCarlo, Zoccolan, & Rust, 2012). We do not claim that ventral stream representations are *sufficient* for visual production, as rendering a physical image still requires translating these features into a motor program to execute the appropriate sequence of actions.

Indeed, future studies could investigate how processes engaged uniquely during visual production but not recognition affect object representations. For example, the role of motor execution could be examined by having participants trace over previously-made drawings, and the role of mental construction could be examined by having participants simulate drawing objects without producing a physical image. The rich sensory feedback generated during production may play an important role in learning, as the visual traces of current and prior movements provide an explicit basis for performance monitoring and error-based updating (Wolpert, Diedrichsen, & Flanagan, 2011; Taylor, Hieber, & Ivry, 2013). In particular, it may be that such sensory feedback is particularly important for learning how to realistically render objects from observation, more so than for informal sketching from semantic knowledge. The functional value of these external traces is important to understand because they are unique to visual production and not shared with other generative cognitive processes known to influence learning, such as selective attention (Chun & Turk-Browne, 2007; Uncapher & Rugg, 2009; Fan & Turk-Browne, 2013) and memory retrieval (Slamecka & Graf, 1978; Crutcher & Healy, 1989; Karpicke & Roediger, 2008).

In the present study, the benefits of training for production and recognition were specific to the objects participants had practiced drawing. However, a hallmark of learning is generalization; thus, an important goal for future research is to understand the training conditions under which participants improve (or worsen) their ability to draw objects they did not practice, a form of skill learning also known as structural learning or learning-to-learn (Braun, Aertsen, Wolpert, & Mehring, 2009; Lake, Salakhutdinov, & Tenenbaum, 2015). Various factors merit detailed investigation, including the amount of training, variability in the objects and categories practiced (Schmidt & Bjork, 1992; Wrisberg & Liu, 1991), the type of performance feedback returned (Taylor et al., 2013; Nikooyan & Ahmed, 2015), and the effects of progressive training sequences from simpler to more complex visual forms, as advocated in classic instruction manuals (Ruskin, 1881).

Humans draw for many reasons: including to depict, to record, to plan, to explain, and to create (Tversky, 2011). Just as investigations of both verbal comprehension and production are indispensable to theories about linguistic communication, a more complete understanding of visual communication will require examining how visual recognition and production interact to support behavioral goals. Ultimately, inquiries into the psychological basis of visual production may shed new light upon the origins of symbolic writing systems for communication, and the nature of our ability to apprehend abstract meanings from visual artifacts.

## Code availability

The code for the analyses presented in this article is publicly available in a Github repository: https://github.com/judithfan/common_reps_production_recognition

## Data availability

The data presented in this article is publicly available in a figshare repository: https://figshare.com/articles/common_repns_production_recognition_data/4503092

## Acknowledgements

This work was supported by NSF GRFP DGE-0646086, NIH R01 EY021755, R01 MH069456, and the David A. Gardner’69 Magic Project at Princeton University. Thanks to Ken Norman, Jordan Suchow, Jordan Taylor, and the Turk-Browne lab for helpful comments. Thanks also to NVIDIA for a grant of GPU hardware, and to Lauren Aleza Kaye for designing the 3D morph stimuli.

## Author contributions statement

J.E.F. and D.L.K.Y. formulated initial idea, performed computational modeling, and analyzed data. J.E.F. performed the experiments. J.E.F., D.L.K.Y., and N.B.T.-B. designed experiments, planned analyses, interpreted results, and wrote the paper.

## Additional information

The authors declare no competing interests.

